# The Lifeact-EGFP Quail: A New Avian Model For Studying Actin Dynamics *In Vivo*

**DOI:** 10.1101/2023.11.19.567639

**Authors:** Yanina D. Alvarez, Marise van der Spuy, Jian Xiong Wang, Ivar Noordstra, Siew Zhuan Tan, Murron Carroll, Alpha S. Yap, Olivier Serralbo, Melanie D. White

## Abstract

Here we report the generation of a transgenic Lifeact–EGFP quail line for the investigation of actin organization and dynamics during morphogenesis *in vivo*. This transgenic avian line allows for the high-resolution visualization of actin structures within the living embryo, from the subcellular filaments that guide cell shape to the supracellular assemblies that coordinate movements across tissues. The unique suitability of avian embryos to live imaging facilitates the investigation of previously intractable processes during embryogenesis. Using high-resolution live imaging approaches, we present the dynamic behaviours and morphologies of cellular protrusions in different tissue contexts. Furthermore, through the integration of live imaging with computational segmentation, we reveal the dynamics of cells undergoing apical constriction and the emergence of large-scale actin structures such as supracellular cables and rosettes within the neuroepithelium. These findings not only enhance our understanding of tissue morphogenesis but also demonstrate the utility of the Lifeact–EGFP transgenic quail as a new model system for live *in vivo* investigations of the actin cytoskeleton.

## Introduction

During morphogenesis, tissue is formed by a complex interplay of changes in cellular geometry and cellular movements. The forces required to shape developing tissues are generated by actomyosin networks that are coupled to cell-cell and cell-matrix junctions (Clarke and Martin, 2021; Perez-Vale and Peifer, 2020). Contractions of the actomyosin network allow cells to pull on their neighbours or the matrix. These active forces at the cellular level are translated into dramatic changes in shape at the tissue level which underly morphogenesis.

Remodelling of the actin cytoskeleton is the basis of a toolbox of common morphogenetic processes that drive tissue formation. These range from the subcellular scale, such as cellular protrusions, to changes in shape at the cellular scale, up to multicellular structures such as actin cables and cellular rosettes. Live imaging of actin dynamics has emerged as a powerful tool to unravel these processes (Blankenship et al., 2006; Christodoulou and Skourides, 2015; Fierro-Gonzalez et al., 2013; Phng et al., 2013; Samarage et al., 2015; Schimmel et al., 2020; Zenker et al., 2018).

To visualise actin dynamics in real-time during morphogenesis in a highly accessible vertebrate model, we generated the TgT2[UbC:Lifeact-EGFP] transgenic quail. This transgenic line, expressing an EGFP fusion of the actin-binding Lifeact peptide (Riedl et al., 2008), allows for the high-resolution visualization of actin structures within the living embryo, from the subcellular filaments that guide cell shape to the supracellular assemblies that coordinate movements across tissues. The unique suitability of avian embryos to live imaging facilitates the investigation of previously intractable processes during embryogenesis (Benazeraf et al., 2010; Huss et al., 2015; Li et al., 2019; Nerurkar et al., 2019; Saadaoui et al., 2020; Xiong et al., 2020). We present novel insights into the dynamic behaviours of cellular protrusions in different tissues, apically constricting cells, and the assembly of multicellular structures. These findings not only enhance our understanding of tissue morphogenesis but also demonstrate the utility of the TgT2[UbC:Lifeact-EGFP] transgenic quail as a new model system for live *in vivo* investigations of the actin cytoskeleton.

## Results and Discussion

### Generation of a transgenic Lifeact-EGFP quail line

To avoid potential overexpression artifacts, we used the low-expression ubiquitin promoter (Qin et al., 2010) to drive Lifeact-EGFP expression in our transgenic quail line (TgT2[UbC:Lifeact-EGFP]). The UbC-Lifeact-EGFP cassette was inserted in between 5’
s and 3’ Tol2 (T2) transposable elements to facilitate stable integration into the quail genome (Fig. 1A). Using a direct injection technique (Barzilai-Tutsch et al., 2022; Serralbo et al., 2020; Tyack et al., 2013) we transfected the blood-circulating primordial germ cells *in vivo*. Fifty wild-type embryos at stage HH16 (E2.5) were injected in the dorsal aorta with a mix of lipofectamine 2000, the UbC-Lifeact-EGFP plasmid and a pCAG-Transposase construct. One male and one female founder were identified and mated with wild-type quails to establish lines. After further breeding the lines were indistinguishable and the line from the male founder was selected for long-term maintenance. The TgT2[UbC:Lifeact-EGFP] quails are viable, phenotypically normal and fertile and can be maintained as heterozygotes or homozygotes. The transgene inheritance followed a Mendelian distribution, indicating that the gene was likely integrated at a single location in the genome. We could detect robust Lifeact-EGFP expression from Hamilton and Hamburger stage 3 (HH3) onwards (Hamburger and Hamilton, 1951) (Figs. 1B, S1). The Lifeact-EGFP fluorescence overlapped with Phalloidin-568 and SPY650 FastAct staining in fixed embryos and was enriched in regions of increased Phalloidin-568 intensity (Figs. 1B - E).

**Figure 1.**
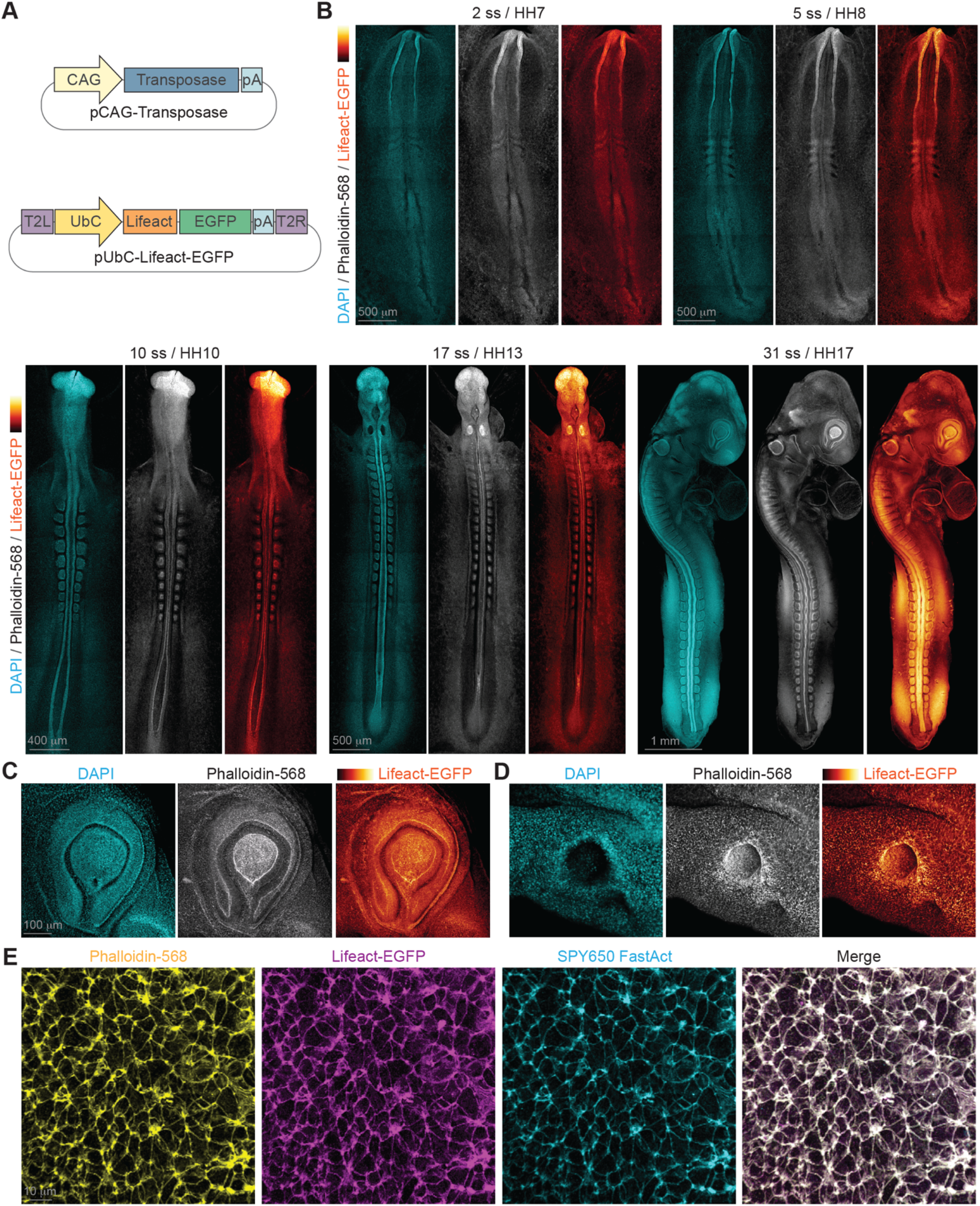
The endogenous actin cytoskeleton is fluorescently labelled in the developing TgT2[UbC:Lifeact-EGFP] transgenic quail embryo. (A) Plasmid vectors used to generate the transgenic line. (B) Tiled confocal images show the Lifeact-EGFP signal matches labelling of the actin cytoskeleton by Phalloidin-568 throughout early embryonic development. Lifeact-EGFP signal is enriched in regions of increased Phalloidin-568 labelling in the eye (C) and the otic pit (D). Higher magnification images demonstrate the similarity of the Lifeact-EGFP signal to staining with Phalloidin-568 and SPY650FastAct.

### Live imaging of actin dynamics during morphogenesis

We performed live imaging of the TgT2[UbC:Lifeact-EGFP] quail line at a range of scales from the subcellular to the tissue level to examine the actin cytoskeleton dynamics during common morphogenetic tissue processes.

### Imaging cellular protrusions

Many actively migrating cells reorganize their actin cytoskeleton and generate polarized protrusions (Schaks et al., 2019). Although this has been visualised in multiple tissues (Lamb et al., 2020; Mishra et al., 2019; Olson and Nechiporuk, 2021; Omelchenko, 2022), the actin cytoskeleton has never been imaged live at high resolution during fusion of the early heart tube *in vivo*. Vertebrate heart formation begins with the migration of bilateral sheets of cardiogenic mesoderm towards the midline where they fuse to form the primitive heart tube (Kirby, 2007; Rosenquist, 1966; Stalsberg and DeHaan, 1969). Here we performed high spatiotemporal resolution live imaging of mesodermal cardiac progenitors crawling over the underlying endoderm of the Anterior Intestinal Portal (AIP) in the TgT2[UbC:Lifeact-EGFP] quail embryo at HH7 – HH8 stage (E1). The bilateral collectives of cardiac progenitor cells form many filopodia which are enriched in Lifeact-EGFP and protrude outwards, making contact with the surrounding tissues (Fig. 2A, Supplementary movie 1). Fusion of the bilateral heart fields is initiated when progenitor cells from each side first make contact via their filopodia (Fig. 2B). Similar to the migration of Drosophila myotubes (Bischoff et al., 2021), we did not find any lamellipodia suggesting the cardiac progenitor cells use a filopodia-dependent migratory mechanism. Cardiac progenitor cell filopodia are on average 9.1mm +/-0.5mm long and highly dynamic with an average persistence time of 389.1 s +/-22.9 s (n = 86 filopodia, 4 embryos). Filopodia that contact the surrounding tissues are significantly longer and more persistent than those that do not make contact (11.2mm +/-0.7mm, n = 42 and 523.6 s +/-34.5 s, n = 37, compared to 7.2mm +/-0.4mm, n = 44 and 276.0 s +/-20.5 s, n = 44, Fig 2C – E). The tissues surrounding the cardiac progenitor cells are covered in an extracellular matrix rich in fibronectin, which also extends along some of the filopodia (Fig. S2). As integrins are known to be present at filopodial tips (Galbraith et al., 2007; Lagarrigue et al., 2015), the higher persistence of filopodia in contact with surrounding tissues may indicate a force-dependent stabilisation of the filopodia (Alieva et al., 2019). This indicates these filopodia could have signalling roles as proposed previously (Francou et al., 2014) and/or mechanical roles during cardiac progenitor cell migration.

**Figure 2.**
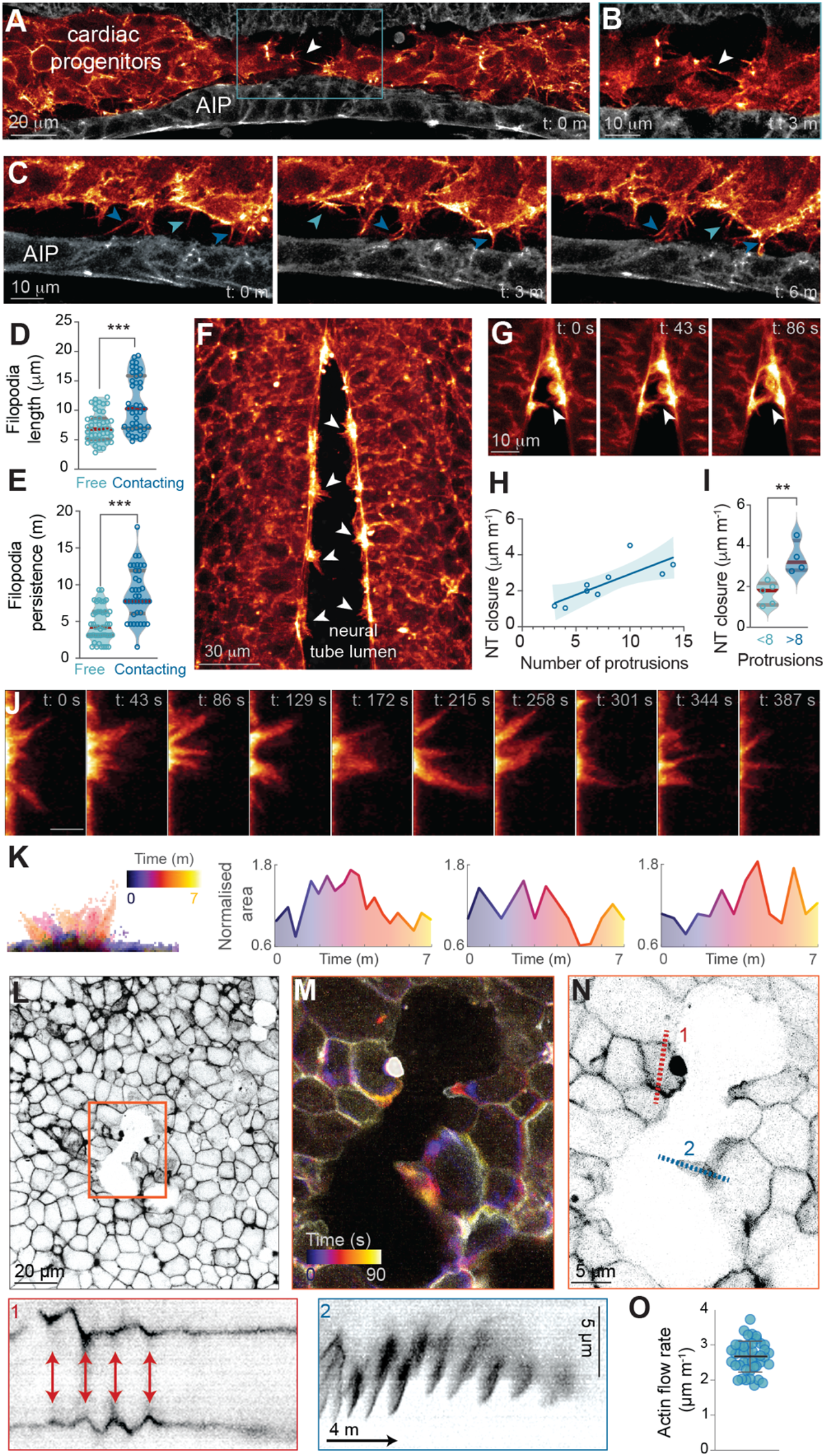
Live imaging reveals dynamics of cellular protrusions in different tissue contexts. (A) Mesodermal cardiac progenitor cells extend filopodia towards the surrounding tissues as the bilateral heart fields migrate towards the midline over the Anterior Intestinal Portal (AIP). White arrowhead indicates leading cardiac progenitor cells extending filopodia towards each other. The tissues surrounding the cardiac progenitor cells are pseudocoloured for clarity. (B) Inset of boxed region in A 3 minutes later shows the first contact via a filopodium (white arrow) between the leading cardiac progenitor cells from each side. (C) Time series of filopodia projecting from cardiac progenitor cells towards the AIP. Filopodia contacting AIP cells (dark blue arrowheads) are longer and have higher persistence than free filopodia (light blue arrowheads). Quantified in D and E, n = 86 filopodia from 4 embryos. *** p < 0.001 (Welch’s test for D and bilateral Mann-Whitney test for E). Violin plots show mean and first and third quartiles. (F) Cellular protrusions of varying morphologies (white arrowheads) extend from the edges of the neural folds into the open lumen of the neural tube. (G) Protrusions reach across the neural tube lumen (white arrowhead) and contact the opposite neural fold close to the zippering point. (H, I) Neural tube (NT) closure is faster in embryos with more protrusions. Shaded area in (H) indicates 95% C.I. **p < 0.01 (Student’s t-test) in (I). Violin plots show mean and first and third quartiles. (J) High spatiotemporal resolution imaging reveals the highly dynamic nature of the neural tube protrusions. (K) Computationally masking the protrusions (left panel) allowed measurement of the area change over time, three examples shown from 2 embryos. L) The ectodermal sheet contains transient apertures indicated by orange box. (M) Higher magnification of inset from L, pseudocoloured with a temporal code to reveal dynamic actin-based structures. (N) Kymographs made from regions indicated demonstrate contractile pulsations in cell junctions (1) and formation of lamellipodia by actin flow (2). (O) Speed of actin flow measured at lamellipodia, n = 39 flow waves from 4 cells. Graph shows mean +/-s.d.

To examine cellular protrusions in a different tissue context we turned to the developing neural tube. Unlike the stereotypical filopodia of the cardiac progenitor cells, cellular protrusions of varying morphologies are proposed to be required for correct neural tube closure (Rolo et al., 2016). Despite many descriptions of these protrusions in fixed specimens as cytoplasmic threads, ruffles, filopodia, blebs or lamellipodia over the last 50 years (Bancroft and Bellairs, 1975; Geelen and Langman, 1979; Rolo et al., 2016; Waterman, 1976), there has been very little live imaging of their dynamics (Massarwa and Niswander, 2013; Pyrgaki et al., 2010; Ray and Niswander, 2016). We performed high spatiotemporal resolution live imaging of TgT2[UbC:Lifeact-EGFP] quail embryos during spinal neural tube closure at stage HH9 – HH10 (E1) (Ainsworth et al., 2009). Clusters of protrusions highly enriched for Lifeact-EGFP were clearly visible protruding into the open lumen of the neural tube (Fig. 2F). Close to the zippering point of the closing neural tube, some protrusions reached across the open neural tube lumen, contacted the opposing neural fold and appeared to assist in pulling the neural folds together (Fig 2G, Supplementary movie 2). Zippering of the neural tube progressed at a faster rate in embryos with more protrusions (Figs. 2H, I), supporting an active role for the protrusions in neural tube closure. The morphology of these protrusions was variable, consisting of both lamellipodia-like and filopodia-like structures (Fig. 2J), which constantly changed shape and area for >60 mins until the approaching zippering point of the neural tube reached them (Figs. 2J, K, Supplementary movie 3). Although these neural tube protrusions have been investigated for decades, live imaging of the TgT2[UbC:Lifeact-EGFP] quail has enabled the first high-resolution visualisation of the dynamics and behaviour of neural tube protrusions, and suggests a potential mechanical role for these structures in zippering of the neural folds.

Building on the observed dynamic protrusions in the developing neural tube, we shifted our focus to the migrating surface ectoderm which is also crucial for neural tube formation (Christodoulou and Skourides, 2022; Galea et al., 2017; Maniou et al., 2021; Marshall et al., 2023; Moury and Schoenwolf, 1995). As the neural tube is closing, the surface ectoderm migrates over it towards the embryonic dorsal midline. During this migration, transient apertures in the ectodermal sheet were observed (Fig. 2L). High-speed spinning disk confocal imaging of the TgT2[UbC:Lifeact-EGFP] quail embryo revealed distinctive oscillations of the Lifeact-EGFP at cell junctions and the formation of lamellipodia as surrounding cells tried to close the gap (Fig. 2M, Supplementary movie 4). The fluctuating cell junctions are indicative of pulsed actomyosin contractions and occur at intervals of approximately 2 minutes (Fig. 2N, box 1). Simultaneously, a retrograde actin flow at an approximate rate of 2.6 mm min^-1^ was identified, propelling the formation of lamellipodia (Figs. 2N, box 2, O). Interestingly, the lamellipodia appeared to form preferentially along the gap edges with the lowest curvature as described previously in vitro (Anon et al., 2012). These lamellipodia, in turn, facilitated the forward movement of cells, contributing to the resolution of the temporary openings and sealing of the ectodermal sheet.

Together, our live imaging confirmed that the TgT2[UbC:Lifeact-EGFP] quail line is an excellent model system for studying the rapid remodelling of the actin cytoskeleton during protrusive cell behaviours *in vivo*.

### Actin dynamics during apical constriction

Another common morphogenetic process during tissue development is apical constriction, in which cells change geometry by shrinking their apical surface (Martin and Goldstein, 2014). Epithelial remodelling requires apical constriction in a wide variety of contexts to awain the correct form and architecture of the tissue. Different dynamic patterns of actomyosin network contraction can drive apical constriction (Martin and Goldstein, 2014). In invertebrates, pulsed contractions of a medioapical meshwork drive shrinkage of the apical cell surface (Martin et al., 2009; Solon et al., 2009). However, a purse-string-like contraction of a circumferential cellular actomyosin cable has long been proposed to drive apical constriction in vertebrates (Baker and Schroeder, 1967; Schroeder, 1970). More recent work demonstrated that pulsatile contraction of a transient apical actin network drives stepwise shrinkage of the apical surface in constricting cells in the Xenopus neuroepithelium (Christodoulou and Skourides, 2015) and the mouse epiblast (Francou et al., 2023), but this has not yet been investigated in the avian neural plate.

To visualise the actin cytoskeleton and cell dynamics during apical constriction we performed live imaging of the neuroepithelium in the open region of the neural tube of TgT2[UbC:Lifeact-EGFP] quail embryos at HH8 – HH9 (E1) (Fig. 3A). Computational segmentation was performed to identify cells undergoing apical constriction. To achieve this, we first used a custom local Z projection strategy to project the curved apical surface of the neuroepithelium onto a 2D plane. Next, we used the Cellpose 2.0 segmentation algorithm to identify cell boundaries labelled by Lifeact-EGFP. Using a custom MATLAB script, we identified cells undergoing apical constriction – defined as cells with a rate of area decrease > 0.02 m^-1^ (Fig. 3B). Measuring the ratio of Lifeact-EGFP signal at the apical cortex relative to the cell junctions revealed an average increase of 71.7%+/-2.9 % during the first 25% of the reduction in apical cell area (Figs. 3C, S3A-B). The inverse correlation between mean Lifeact-EGFP intensity at the apical cortex and mean apical cell area is highly significant (Fig. S3B). Furthermore, the identified cells did not undergo a constant decrease in apical cell area but instead showed a more pulsatile pattern consistent with a ratchet-like mechanism (Figs. 3C, D). There was a moderate, but highly significant correlation between the rate of change in Lifeact-EGFP intensity at the apical cortex and the change in apical cell area for individual cells (Fig. S3C). Pulsatile ratchet-like apical constrictions have been well-studied in Drosophila where they drive rapid apical constriction en masse during gastrulation and dorsal closure (Martin et al., 2009; Mason et al., 2013; Solon et al., 2009). However, the apical constriction events identified here are more similar to those recently described during epiblast cell ingression at the mouse primitive streak where isolated cells constrict their apical surface over a 25 – 90 minute period (Francou et al., 2023). Although the apical surface area of all cells we tracked in the quail neuroepithelium fluctuated over time, these area fluctuations were significantly larger in cells undergoing apical constriction (Figs. 3D, E). Consistent with observations in the mouse primitive streak, the rate of area change in apically constricting quail neuroepithelium cells was faster during constriction phases than expansion phases (Fig. 3F). Our findings support the growing evidence that apical constriction driven by a ratchet-like mechanism is a common process shared by invertebrates and vertebrates.

**Figure 3.**
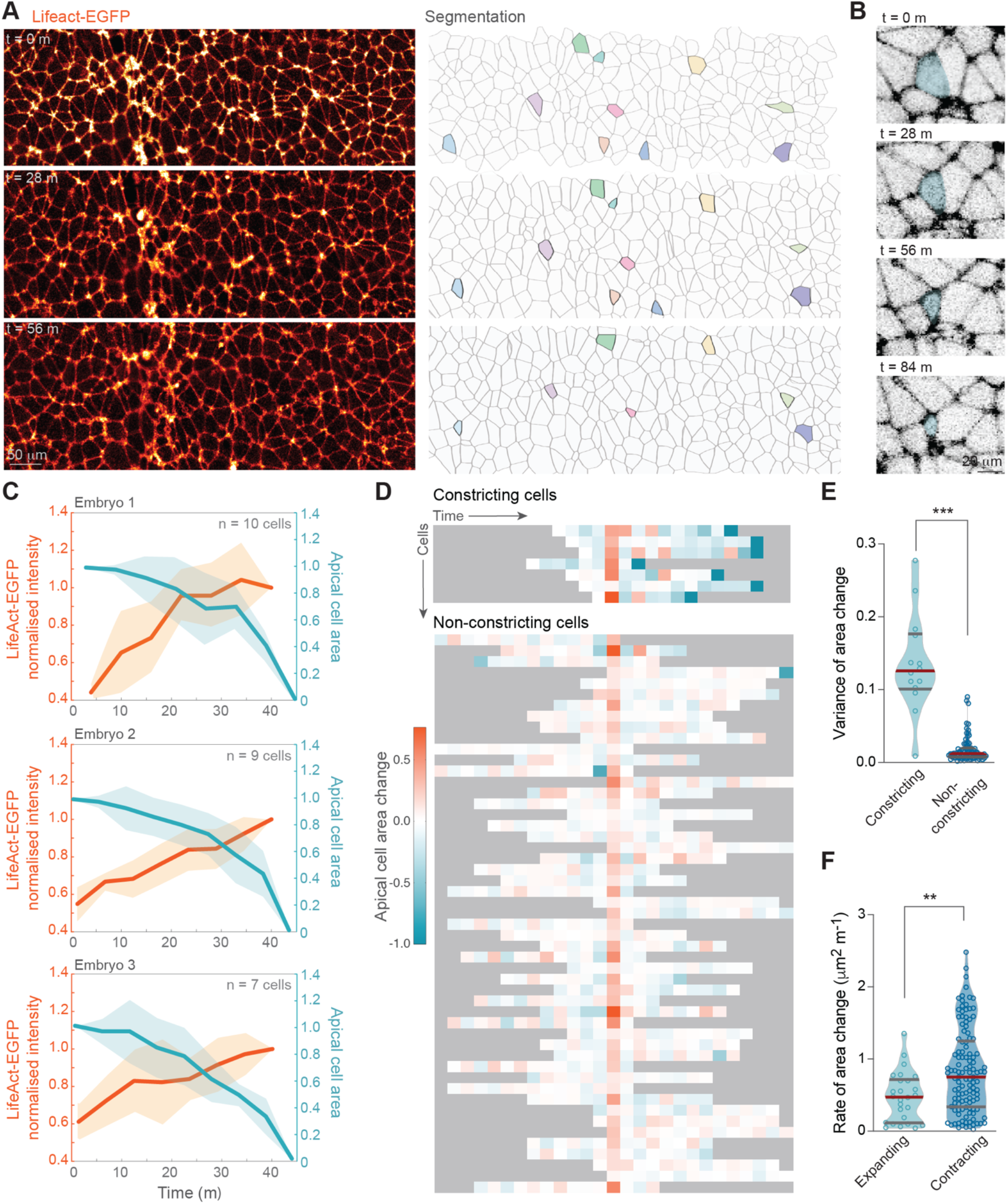
Cells undergo pulsatile apical constriction in the developing neuroepithelium. (A) Computational segmentation (right panel) of the Lifeact-EGFP signal (ler panel) in live imaging of the neuroepithelium identifies cells undergoing apical constriction. (B) Time series of an apically constricting cell (blue cell in segmented image in A). (C) The ratio of Lifeact-EGFP intensity at the apical cortex relative to the cell junctions increases in a pulsatile manner as the apical cell area shrinks. Graphs show mean (coloured line) +/-s.e.m (shading) for the normalised Lifeact-EGFP intensity and apical cell area over time in 3 embryos. (D) Heatmap of apical area change over time in cells undergoing apical constriction and non-constricting cells. Example from 1 embryo, n = 7 constricting cells and 47 non-constricting cells. Cells are aligned by the maximal observed area. (E) Constricting cells have a higher variance of apical area change than non-constricting cells, n = 14 constricting cells and 62 non-constricting cells from 2 embryos. *** p < 0.001 (bilateral Mann-Whitney test). (F) Rate of area change during expansion and contraction phases of constricting cells, n = 19 cells from 3 embryos. Area change is faster during contraction phases. **p < 0.01 (bilateral Mann-Whitney test). Violin plots show mean and first and third quartiles.

### Formation of supracellular actin cables and multicellular rosettes

The formation of supracellular actin cables and multicellular rosettes is associated with tissue reorganization in a variety of species (Harding et al., 2014; Miao and Blankenship, 2020). Supracellular actin cables mechanically link multiple cells to allow the coordinated application of tension across the tissue. They can also reinforce the structure of entire tissues and increase their resistance to mechanical strain (Duda et al., 2019). Myosin-driven contraction of supracellular actin cables constricts the lateral edges of the cells along the cable, pulling them into a rosette structure with a centrally shared interface. These multicellular rosettes are a common feature during morphogenesis of various organs (Gompel et al., 2001; Lienkamp et al., 2012; Villasenor et al., 2010).

In the avian embryo, supracellular actin cables and epithelial rosettes are thought to be important for the formation of the neural tube (Nishimura et al., 2012; Nishimura and Takeichi, 2008). The planar polarized assembly of supracellular actin cables is proposed to enable polarized tissue contraction along the mediolateral axis. This is thought to allow the tissue to bend only along the anteroposterior axis of the embryo, rather than bending radially if the contraction was isotropic (Nishimura et al., 2012). However, this idea is based on computational modelling and studies on fixed tissue which is prone to shape artifacts arising from the fixation. Using live imaging, we visualised the formation of supracellular actin cables and rosettes in the neuroepithelium of the TgT2[UbC:Lifeact-EGFP] quail embryo at HH8 – HH9 (E1) (Fig. 4A). We focused on the region caudal to the closed neural tube, where the neural plate was still relatively flat. The transverse view of the 3D stack revealed a slight curvature along the midline of the neural plate (Fig. 4A, t = 0 h, x,y view). We identified supracellular actin cables spanning 10 or more cells in a mediolateral direction. The actin cables are enriched for di-phosphorylated myosin light chain, suggesting they are contractile (Fig S4A). Live imaging for two hours revealed that the supracellular cables contracted and the curvature of the neural plate increased along the mediolateral axis (Figs. 4A, t = 2 h, x,y view, B).

**Figure 4.**
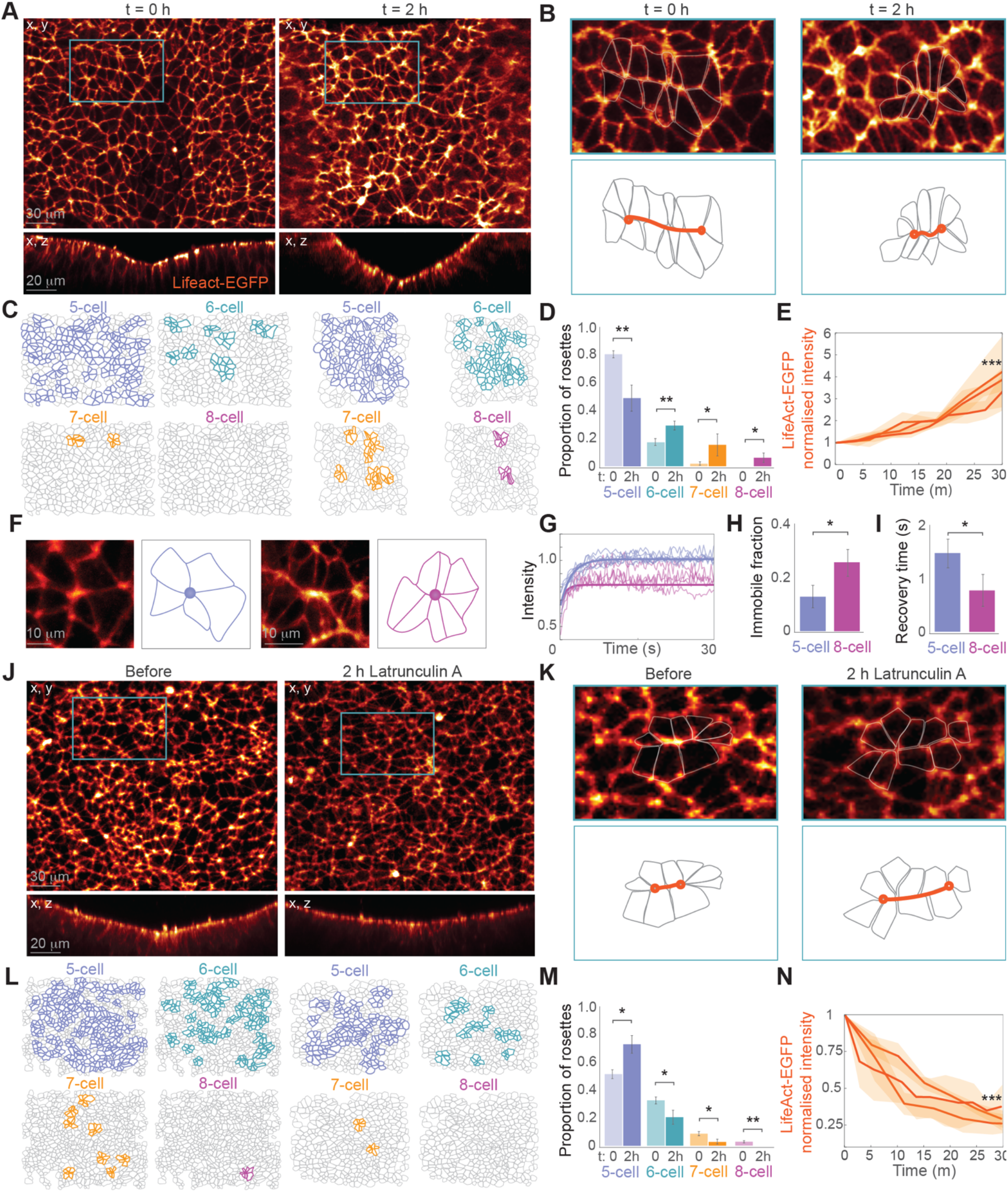
Formation of supracellular actin cables and multicellular rosettes in the developing neuroepithelium. (A) Supracellular actin cables (blue boxes) form in the relatively flat neural plate and contract within 2 hours, concomitant with bending along the midline (x,z views). (B) Insets show supracellular cables indicated in A. (C) Computational segmentation of areas shown in A reveals an increasing number and complexity of rosettes over time, quantified in (D). Bar graphs show mean +/-s.e.m, n = 827 rosettes from 4 embryos. **p<0.01, **p<0.05, Student’s t-test. (E) The intensity of Lifeact-EGFP increases as the supracellular cable contracts (normalised to cable area). Graph shows mean (orange lines) +/-s.e.m (shaded areas) from 3 embryos. ***p<0.001, Wilcoxon Signed Rank Test, t: 0m vs t: 30m. (F – I) FRAP of the actin at the central vertex shows a higher immobile fraction and a faster recovery time in 8-cell rosettes compared to 5-cell rosettes. Graphs show mean +/-s.e.m. *p<0.05, Student’s t-test (J, K) Treatment with Latrunculin A causes extension of supracellular actin cables and flattening of the neural plate. (L, M) Computational segmentation of areas shown in J reveals that Latrunculin A reduces the number and complexity of rosettes. Bar graphs show mean +/-s.e.m, n = 2044 rosettes from 4 embryos. (N) The intensity of Lifeact-EGFP localised to the actin cable is decreased by Latrunculin A treatment (normalised to cable area). Graph shows mean (orange lines) +/-s.e.m (shaded areas) from 3 embryos. ***p<0.001, Wilcoxon Signed Rank Test, t: 0m vs t: 30m.

As contraction of supracellular actin cables can generate multicellular rosettes, we used computational segmentation and a custom MATLAB script to identify rosettes consisting of 5 – 8 cells (Fig. 4C). The rosettes were enriched for di-phosphorylated myosin light chain at their central vertex (Fig. S4B). Surprisingly, even when the neural plate was only slightly curved, most cells were already part of a multicellular rosette. 79.8% of these rosettes contained 5 cells, 17.8% contained 6 cells and only 2.9% contained 7 cells (Figs. 4C, D). After two hours, the total number of rosettes had increased by 45.4%. Furthermore, the distribution had shifted towards higher-order rosettes with only 49.4% containing 5 cells but 29.2% containing 6 cells, 8.9% containing 7 cells and 3.7% containing 8 cells. As the actin cables forming the rosettes contracted, the Lifeact-EGFP intensity (normalised to cable area) significantly increased, suggesting continued actin accumulation during cable contraction (Fig. 4E).

To examine the stability of the actin remaining at the centre of the multicellular rosettes following contraction of the supracellular cables we used Fluorescence Recovery After Photobleaching (FRAP). Comparing 5-cell rosettes to 8-cell rosettes revealed a higher immobile fraction and a decreased recovery time in the higher-order rosettes (Figs. 4F – I). Our results show that a greater proportion of the actin molecules are stationary and not exchanging with the surrounding pool of actin within the centre of the 8-cell rosettes. This may indicate a higher degree of actin stabilization or stronger binding interactions that prevent the actin molecules from diffusing away. However, within the mobile pool of actin, there is also a higher rate of actin turnover or a more efficient replenishment in the centre of the higher-order rosettes, possibly due to greater accessibility or a higher rate of actin polymerisation at the site. Together, these findings suggest that the actin structures linking the cells in the centre of the higher-order rosettes are rapidly remodelled, but once actin monomers are integrated into the structure, they become part of a stable network that resists further exchange. These complex dynamics may reflect tuning of the actin remodelling within multi-cellular rosettes to balance structural integrity with the flexibility required for ongoing morphogenesis.

To perturb the supracellular actin cables and rosettes, we treated the embryos with the actin polymerisation inhibitor Latrunculin A (Coue et al., 1987). During two hours of live imaging, we observed a decrease in mediolateral curvature of the neural plate and extension of the supracellular actin cables (Figs. 4J, K). Latrunculin A treatment also decreased the total number of rosettes by 48.0% (Fig. 4L). Moreover, there was a corresponding shift from higher-order towards lower-order multicellular rosettes (Figs. 4L, M). During Latrunculin A treatment there was a significant decrease in the Lifeact-EGFP intensity in the supracellular cables, consistent with a reduction in actin polymerisation (Fig. 4N). Together, our results support a role for supracellular actin cable contraction and rosette formation in the anisotropic bending of the neural plate during neural tube formation. The increasing number and complexity of the multicellular rosettes during neural plate bending suggests that the persistence of these structures is higher than the transitory rosettes involved in tissue remodelling in Drosophila (Blankenship et al., 2006). The accumulation of higher-order rosettes in the neural plate may also contribute to the reported decrease in tissue fluidity during neurulation (Bocanegra-Moreno et al., 2023; Yan and Bi, 2019).

By employing cutting-edge live imaging of a new Lifeact-EGFP transgenic quail model, we have investigated the dynamics of the actin cytoskeleton across various tissues *in vivo*. Our approach has allowed us to observe cellular protrusions and dynamic behaviours like apical constriction and the formation of complex actin-dependent structures, including supracellular actin cables and mulgcellular rosettes, in the developing embryo. The insights gained from this research offer a deeper understanding of processes involved in tissue morphogenesis and confirm the TgT2[UbC:Lifeact-EGFP] transgenic quail as an invaluable resource for real-time, *in vivo* study of the actin cytoskeleton.

## Materials and methods

### Generation of transgenic Lifeact-EGFP quail line

The direct injection technique was performed as described previously (Serralbo et al., 2020; Tyack et al., 2013). Briefly, plasmids were purified using a Nucleobond Xtra Midi EF kit. 1 μl of injection mix containing 0.6 μg of pUbC-Lifeact-EGFP Tol2 plasmid, 1.2 μg of CAG Transposase plasmid and 3 μl of Lipofectamine 2000 CD (ThermoFisher Scientific) in 90 μl of OptiPro was injected in the dorsal aorta of 2.5-day-old embryos. Eggs were then sealed and incubated until hatching. Chicks were grown for 6 weeks until they reached sexual maturity. Semen from the male was collected as described previously (Serralbo et al., 2020). Genomic DNA from semen was extracted and PCR was performed to test for the presence of the transgene. Males showing a positive band were kept and crossed with wild-type females. F1 offspring were selected using GFP goggles and confirmed by genotyping 5 days after hatching by plucking a feather.

### Maintenance of quails

Transgenic quails were hosted and bred at the University of Queensland according to local animal ethical policies. Fertilized quail eggs from transgenic quail lines were collected daily by the animal facility. Eggs were kept at 14°C before use.

### Embryo staining

Embryos were fixed at room temperature for 30 minutes in 4% paraformaldehyde (Sigma Aldrich, P6148-500G), and extra-embryonic tissues were trimmed before adding Phalloidin-Rhodamine (1:1000, Abcam, AB235138), Spy650-FastAct(1:1000, SpiroChrome, SC505) and DAPI (1:1000, Sigma Aldrich, MBD0015-15ML) for 3 hours at room temperature. After incubation, embryos were washed and mounted with fluoromount aqueous mounting medium (Sigma Aldrich, F4680-25ML) for imaging on glass slides (Epredia).

### Whole-mount immunostaining

Embryos were fixed at room temperature for 30 minutes in 4% paraformaldehyde (Sigma Aldrich, P6148-500G), and extra-embryonic tissues were trimmed before immunostaining. Embryos were permeabilised in PBS with 0.5% Triton X-100 (PBTX, Sigma Aldrich, T9284-500mL), and blocked in 0.5% PBTX, 0.2% Bovine Serum (BSA, Sigma Aldrich, A2153-50G) and 0.02% Sodium Dodecyl Sulfate (SDS, Sigma Aldrich, L4509-25G). Embryos were then incubated in primary antibody: Mouse anti-Fibronectin (1:1000, DSHB, B3/D6) or Rabbit anti-Diphospho-MLC (Thr18/Ser19) (1:100, Cell Signalling Technology, 95777S) in blocking buffer overnight at 4°C, followed by washing and incubation in Goat anti-mouse IgG Alexa 647 (1:1000, Thermo Fisher Scientific, A31571) or Goat anti-rabbit Ig Alexa 647 (1:1000, Thermo Fisher Scientific, A21245) for 3 hours at room temperature. After incubation, embryos were washed and mounted on a 6-well glass bottom plate (Cellvis, P06-1.5H-N) with fluoromount aqueous mounting medium (Sigma Aldrich, F4680-25ML) for imaging.

### Quail embryo culture

Eggs were incubated horizontally at 37.5°C in a humidified atmosphere until the desired stages. After incubation, quail eggs were cooled down for approximately 1 hour at room temperature (RT) before culture. While keeping the egg horizontal by placement onto a customized egg holder, 1 – 2 mL of thin albumin was aspirated out with a 5 mL syringe (Nipro) and a 18G sterile needle (BD PrecisionGlide) from the blunt end of the egg to create a space between the embryo and the shell. A small window (∼1.0 × 0.5 cm) was cut on the top of the eggshell to facilitate visualisation and staging. Embryos were staged based on morphological criteria as described previously (Ainsworth et al., 2009). Embryos were cultured according to a protocol described elsewhere (Williams and Sauka-Spengler, 2021). Briefly, the egg yolk and albumin were carefully slid out of the shell into a 35 mm petri dish with the embryo resting at the top of the yolk. The albumin covering the surface of the embryo was carefully wiped away with Kimwipe tissue. A piece of 1.5 × 1.5 cm square filter paper with a 0.2 × 1.0 cm hole in the centre was placed onto the surface of the embryo, with the embryo centered in the hole. The filter paper was then cut along the edge with a pair of sharp dissection scissors to release the embryo from surrounding tissues while maintaining biological tension. The filter paper, with the embryo attached, was lifted away from the yolk using fine forceps and placed onto a 35 mm 1 well dish pre-coated with 1 mL agar-albumin. The yolk was washed away from the embryo using preheated Hanks’ Balanced Salt solution (HBSS). After washing, the embryo was transferred into another 35 mm 1 well dish pre-coated with 1 mL agar-albumin mixture consisting of 0.3% w/v bacto-agar and 50% v/v albumin. Before further use, the 1 well dish containing the embryo was kept in a humidified environmental chamber at RT.

### Latrunculin A treatment

A 20ml drop of 20mM Latrunculin A (Sigma Aldrich, L5163) was placed on the agar-albumin coated plate. Cultured embryos at HH8 – HH9 (E1) were placed on top of the drop dorsal side down, and an additional 20ml drop of Latrunculin A was added to the ventral side. Embryos were allowed to rest in a humidified environmental chamber at 37.5°C for 1 h before imaging.

### Live imaging

Following culture, embryos were allowed to rest for at least 1 h at RT before imaging and then transferred dorsal side facing down to a 6-well imaging plate pre-coated with 250 μL of agar-albumin. All live imaging was done at the IMB microscopy facility (University of Queensland) which is supported by the Australian Cancer Research Foundation.

Live imaging was performed on the following systems: a Zeiss Axiovert 200 Inverted Microscope Stand with LSM 710 Meta Confocal Scanner fitted with dedicated GaAsP 488 nm and 561 nm detectors for increased sensitivity, a Zeiss Axiovert 200 Inverted Microscope Stand with LSM 710 Meta Confocal Scanner, including spectral detection and Airyscan super-resolution detector and Mai Tai eHP 760-1040nm laser with dedicated GaAsP NDD detectors or an Andor Dragonfly spinning disc confocal equipped with dual Andor Zyla 4.2 sCMOS cameras, controlled by Fusion Software (Andor). Microscope environmental chambers were maintained at 37.5°C while imaging.

### Image processing

Images were processed in Imaris 10.0.1 (Bitplane) or FIJI. Pseudocolouring for Figure 2 was performed in Photoshop 2022 (Adobe) by manually selecting and desaturating surrounding tissue areas. Temporal colour coding was performed using FIJI.

### Filopodia quantifications

Length and persistence time was measured in FIJI. The maximal length of a filopodium was measured using a straight segmented line. Persistence was calculated as the final timepoint at which a filopodium is visible minus the first timepoint at which it is observed.

Protrusion dynamics were quantified by the area of the positive pixels in a masked image of the Lifeact-EGFP channel over time. Masking was performed using Otsu’s binarize function in MATLAB.

### Speed of neural tube closure

The closure speed of the neural tube was calculated by tracking the displacement of the zippering point of the neural folds divided by the elapsed time.

### FRAP

FRAP experiments were performed with a Zeiss 40X (W-Plan Apochromat NA1.10) objective at 5-times magnification. A 4 μm x 4 μm region of interest (ROI) was photobleached with 2-photon laser (Mai Tai Laser System) at 840 nm with 30% laser power and imaged every 0.2 seconds for 30 seconds.

For FRAP analysis, mean fluorescence intensity at the photobleached was corrected by background fluorescence and normalized to a non-photobleached reference. The average of the pre-bleach fluorescence intensities was set to 100%. The normalized mean fluorescence intensities were then fitted with an exponential function, to obtain the decay time and the plateau intensity (Ip). The immobile fraction was calculated as I-Ip. The analysis was performed in MATLAB (MATLAB ver. R2021a).

### Lamellipodia dynamics

The ectoderm of the TgT2[UbC-Lifeact-EGFP] transgenic quail at HH7 – 8 was imaged using the Andor Dragonfly spinning disc confocal with a 60x objective and a time interval of 5 seconds. Analysis of lamellipodia dynamics was performed as described previously (Noordstra et al., 2023).

### Computaitonal segmentation

Images were segmented using Cellpose-human in loop workflow (Pachitariu and Stringer, 2022). Cell tracking and rosette detection were performed using custom scripts in MATLAB.

### Detection of apical constriction

All cells were tracked for 50 – 90 minutes. The cell boundaries were converted to polygons in MATLAB and the apical cell area was measured using the polyshape toolkit. Constriction events were identified by normalized apical area rate decrease > 0.02 m^-1^. Rate of area change for each time point was calculated as (area_t+1 – area_t)/ Δ_t. Cell expansion was defined as posigve area rate change and cell constriction was defined as negagve area rate change. The rate of apical area change was defined as: Δarea(t) = (area_t+1 – area_t)/area_t. The variance of area change was calculated for each cell time series.

### Lifeact-EGFP measurements during apical constriction

The intensity of apical surface Lifeact-EGFP was measured by isotropically reducing the segmented cell polygon by 20%. The Lifeact-EGFP at the cell boundary was measured by isotropically expanding the segmented cell polygon by 20% and then subtracting the apical surface measurement.

### Rosette detection

The segmented cell polygons were smoothed using reducepoly function in MATLAB to obtain the positions of cellular vertices. Rosettes were defined as a group of cells with a shared vertex. Shared vertices were identified as those positioned within the sqrt of the mean area of the group of cells.

### Supracellular actin cable measurements

Lifeact-EGFP quangfication during actin cable formation and after Latrunculin treatment was measured with FIJI. The junctions shared by the cells forming the cable were enclosed using the freehand selection tool at each timepoint to measure the mean value of the Lifeact-EGFP channel. The values were normalized to cable area.

### Statistical analysis

Statistical analysis was performed using Prism Graphpad software (version 10.0.2). For all the statistical analysis: ns, p-value > 0.05; *, p-value < 0.05; **, p-value < 0.01; ***, p-value < 0.001.

Correlation coefficients (r) between the mean apical cell area and the mean medial/junctional Lifeact-EGFP intensity for each embryo were obtained using Spearman’s correlation. The same method was used to test the correlation between the change in area and the change in Lifeact-EGFP intensity (medial/junctional) for individual cell over time.

## Acknowledgements

Research was conducted in the Institute for Molecular Bioscience Cancer Biology Imaging Facility, which was established with the support of the ACRF. We thank the IMB Microscopy facility staff.

## Data availability

Data availability: All relevant data can be found within the article and its supplementary information.

## Supplementary Figures

**Figure S1.**
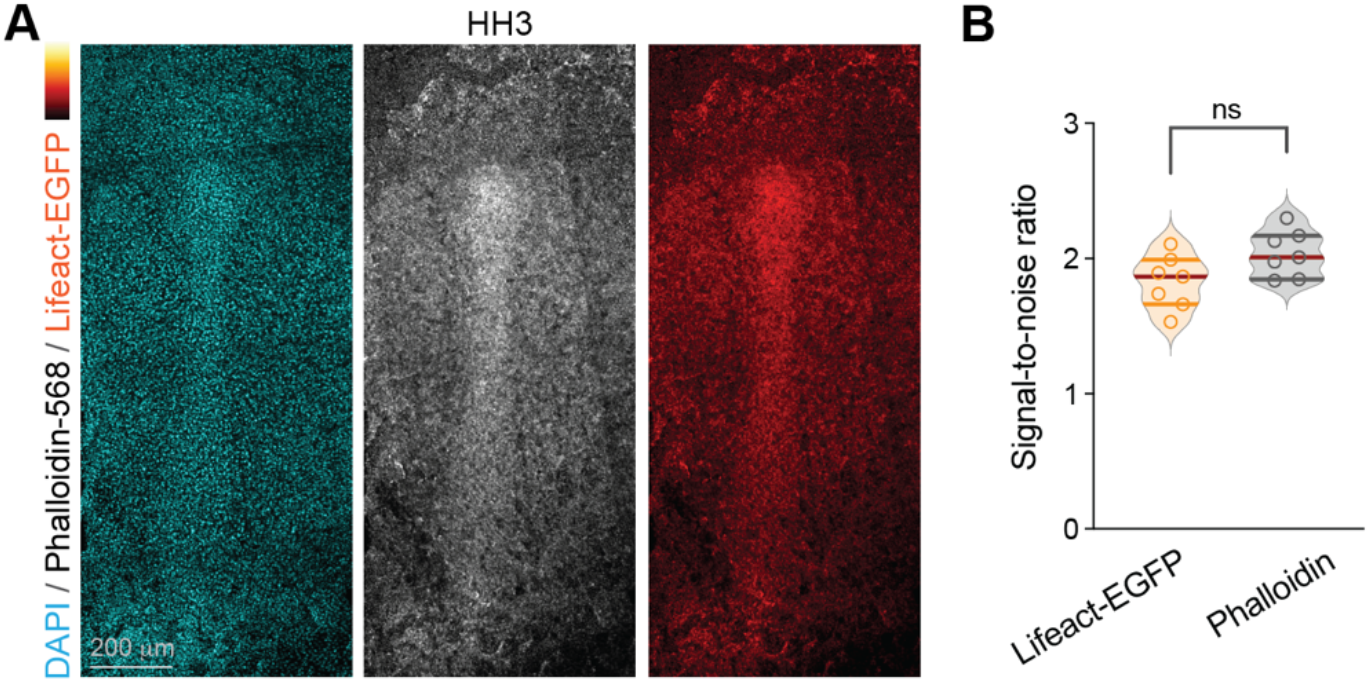
Lifeact-EGFP is expressed at early embryonic stages and has comparable signal-to-noise to Phalloidin. (A) The Lifeact-EGFP signal matches Phalloidin labelling of the actin cytoskeleton in the primitive streak at HH3. (B) There is no significant difference in the signal-to-noise of the Lifeact-EGFP and Phalloidin staining of fixed embryos in Figure 1.

**Figure S2.**
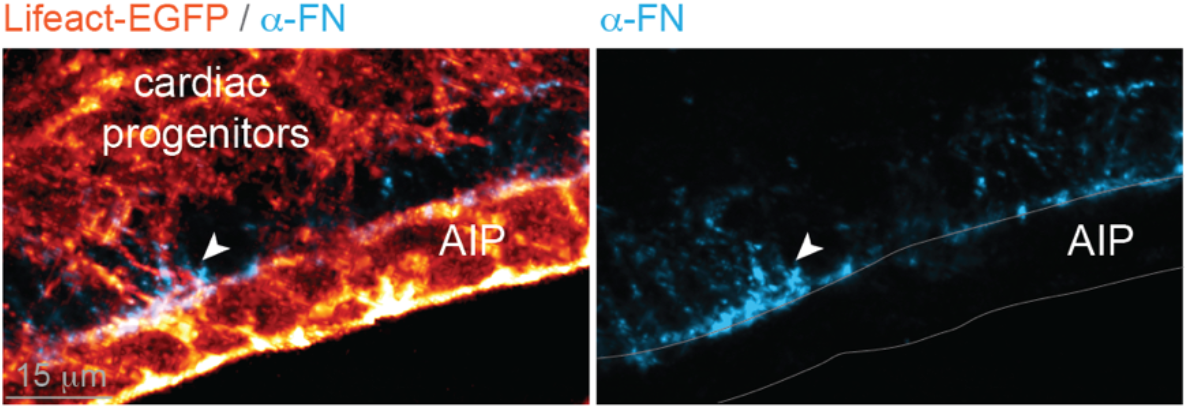
Fibronectin is deposited on the surface of the Anterior Intestinal Portal. Immunostaining shows fibronectin (blue) on the surface of the Anterior Intestinal Portal. (AIP) where filopodia from migrating cardiac progenitor cells make contact (arrowhead). AIP has been outlined for clarity in the right panel.

**Figure S3.**
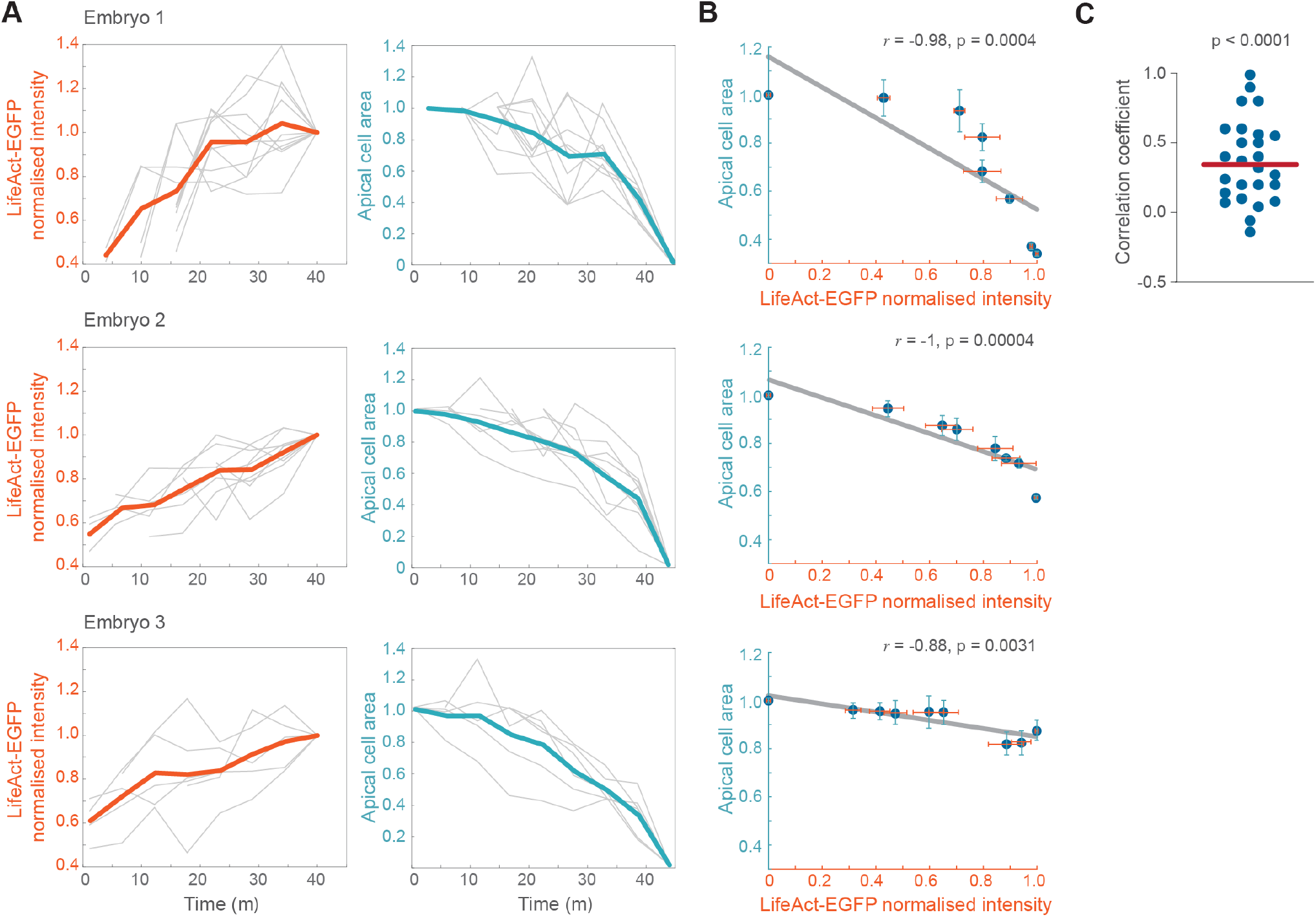
The intensity of Lifeact-EGFP at the apical cortex is inversely correlated with apical cell area. (A) The ratio of Lifeact-EGFP intensity at the apical cortex relative to the cell junctions increases in a pulsatile manner as the apical cell area shrinks. Graphs show mean (coloured line) and individual cells (grey lines) for the normalised Lifeact-EGFP intensity and apical cell area over time in 3 embryos. (B) There is a significant inverse correlation between mean Lifeact-EGFP intensity at the apical cortex (medial/junctional) and mean apical cell area for each embryo. (C) Correlation coefficients for the change in Lifeact-EGFP intensity at the apical cortex versus the change in apical cell area for individual cells. There is a moderate, but highly significant correlation (n = 26 cells from 3 embryos).

**Figure S4.**
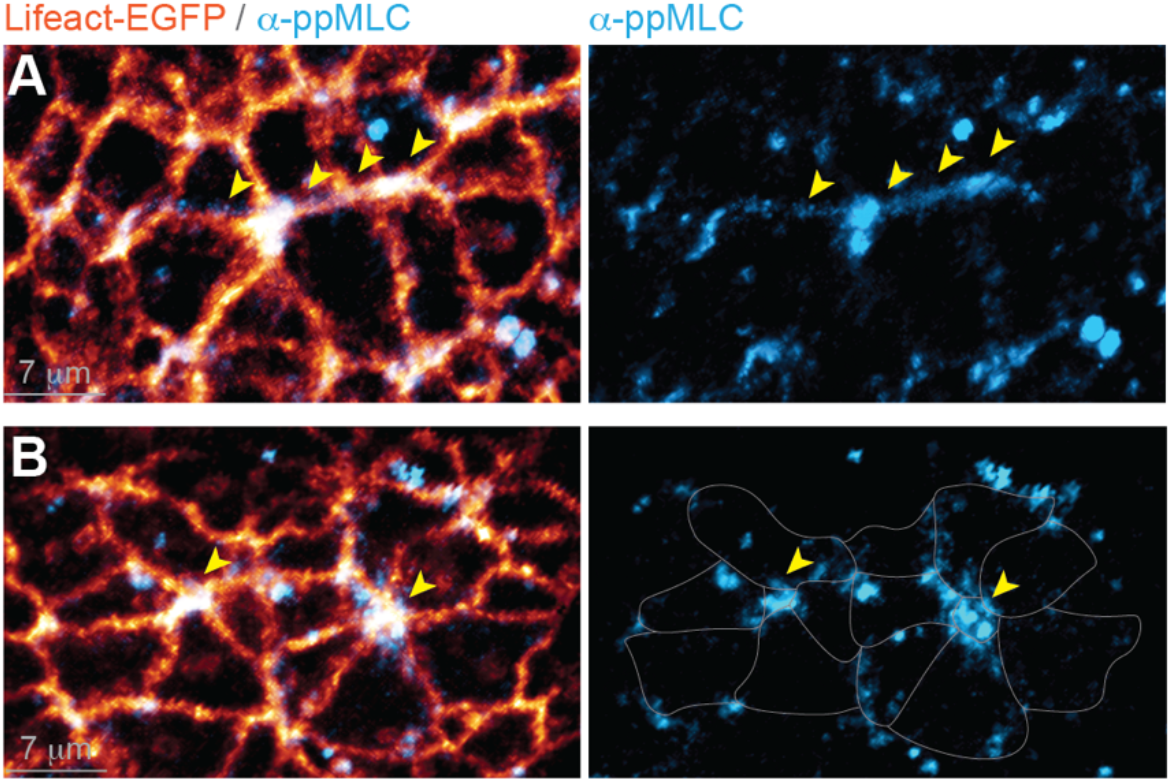
Supracellular actin cables and rosettes are contractile structures. (A, B) Immunostaining reveals di-phosphorylated myosin light chain (blue) is enriched along the supracellular actin cables (arrowheads) and in the central shared vertices of multicellular rosettes. The rosettes have been outlined for clarity in the right panel of B.

**Supplementary Movie 1** Live imaging of mesodermal cardiac progenitors crawling over the Anterior Intestinal Portal in the TgT2[UbC:Lifeact-EGFP] quail embryo at HH7 – HH8 stage (E1). Z-stack images were taken at 40x every 93 seconds. Mesodermal cardiac progenitor cells extend filopodia towards the surrounding tissues as the bilateral heart fields migrate towards the midline. The tissues surrounding the cardiac progenitor cells are pseudocoloured in the last frame for clarity.

**Supplementary Movie 2** Live imaging of cellular protrusions during spinal neural tube closure in a TgT2[UbC:Lifeact-EGFP] quail embryo at stage HH9 – HH10 (E1). Protrusions close to the zippering point reach across the lumen and contact the opposing neural fold. Z-stack images were taken at 40x every 43.5 seconds.

**Supplementary Movie 3** Live imaging of cellular protrusions during spinal neural tube closure in a TgT2[UbC:Lifeact-EGFP] quail embryo at stage HH9 – HH10 (E1). Z-stack images were taken at 40x every 43.5 seconds.

**Supplementary Movie 4** High-speed spinning disk confocal imaging of the ectoderm of the TgT2[UbC:Lifeact-EGFP] quail embryo at HH8 – HH9. Images were taken at 60x every 5 seconds.

**Supplementary Movie 5** Live imaging of the neuroepithelium of the TgT2[UbC:Lifeact-EGFP] quail embryo at HH8-HH9. Z-stack images were taken at 63x every 5.5 minutes. Apically constricting cells are highlighted by computational segmentation in blue.

**Supplementary Movie 6** Live imaging of the neuroepithelium of the TgT2[UbC:Lifeact-EGFP] quail embryo at HH8-HH9. Z-stack images were taken at 63x every 5.5 minutes. Computationally segmented cells (purple) are forming a multicellular rosette.

